# A broken power-law model of heart rate variability spectra in sleep

**DOI:** 10.1101/2025.11.01.685631

**Authors:** Bence Schneider, Martin Dresler, Ferenc Gombos, Ilona Kovács, Róbert Bódizs

## Abstract

**Aims:** The aim of the study was to introduce a parametric description of RR-interval spectra using a broken power-law model, in addition to the parametrization of oscillatory peaks. Furthermore, to use this model to evaluate effects of age, sex and sleep architecture on overnight heart rate variability (HRV) in healthy subjects.

**Methods & Results:** From a polysomnography database, 215 whole-night, high quality electrocardiograms (ECGs) were extracted. The fractal and oscillatory power-spectral densities (PSDs) were calculated from evenly resampled RR-interval time-series, then a broken power-law model was fitted using piecewise linear regression to the double-logarithmic PSD, determining a custom breaking point in the fractal component, and allowing for two independent spectral slopes in the lower and higher frequency domains. The two-slope model provided a more optimal description compared to linear regression in all cases, even when penalizing increased model complexity. Peak detection was applied to the oscillatory component in the LF (0.04–0.15 Hz) and HF (0.15–0.4 Hz) bands, extracting the frequency and prominence of the dominant peak from each.

The high frequency domain intercept, the breaking point frequency and the LF peak frequency decreased significantly with age. Both slopes were flatter in females, while the high domain intercept and the HF peak prominence was significantly increased. Waking after sleep onset and lightest sleep (N1) were associated with lower intercept values, while REM sleep had an opposite effect.

**Conclusions:** The broken power-law model proved to be more appropriate for the description of RR-interval spectra than the single-slope model, and also captured effects of age, sex and sleep structure that were corroborated by the literature. We would like to highlight that while HRV changes are often assumed to be of oscillatory origin, the fractal component has a major contribution to the total PSD, and thus to all measures derived from it.

## Introduction

Biological rhythms in the human body are generally complex as they are a result of multi-level interactions ranging several spatial and temporal scales. The modulation of the heart rate is no exception, being influenced by molecular factors on the millisecond scale, autonomic nervous regulation from seconds to minutes, endocrine effects at the minute to hour rate, and variations in gene transcription on the order of hours/days. Besides endogenous effects, the healthy heart rate constantly adapts to externally determined physical conditions and psychological cues.[Monfredi & Lakatta, 2019]. The degree of this variability can be quantified by heart rate variability (HRV) indices, which express the beat-to-beat time interval variance over a given period, and can be measured by RR-interval changes in the electrocardiogram (ECG). In addition to cardiovascular diseases and autonomic neuropathies, altered HRV values are observed in numerous psychopathologies, like depression, anxiety disorders, bipolar disorder, ADHD, schizophrenia, and mild cognitive impairment [ChuDuc et al. 2013].

Sleep is a state during which most external stimuli are ignored, as a consequence it provides a window on endogenous modulations of different physiological systems, a prominent example being the processes in the autonomic nervous system (ANS). Besides cardiovascular regulation, both the sympathetic and vagal branches of the ANS play an important role in sleep initiation and homeostasis, forming a bidirectional link between heart and sleep dynamics, thus HRV is an indicator of cardiac and sleep health alike. Heart rate variability is atypical in primary sleep disorders like insomnia, or sleep disordered breathing diseases, the most common being obstructive sleep apnea [Tobaldini et al. 2013] At the same time, improper sleep is known to contribute to a range of cardiovascular diseases [Nagai et al. 2010].

Apart from pathologies, significant changes in the ANS also occur with aging, affecting many systems, manifested as alterations in blood pressure, heart rate, breathing, digestion and sleep. With the dysregulation of the ANS, the adaptivity of the system is reduced, which is also reflected by decreased HRV values [Alrosan et al. 2024]

Beyond the fluctuation of interbeat intervals expressed by the time-domain HRV indices, it is possible to gain information about the contribution of different frequencies to this total variance by computing the power-spectral density (PSD) of RR-interval series. Examining the RR-interval power spectra, characteristic oscillatory peaks have been identified and the so-called low frequency 0.04–0.15 Hz (LF) and high frequency 0.15–0.4 Hz (HF) bands were established in the frequency domain HRV methodology, as these bands usually contain the oscillations corresponding to Mayer waves (∼0.1 Hz) and to respiratory sinus arrhythmia (∼0.2 Hz), their increased power have been conventionally attributed to sympathetic and parasympathetic cardiac nerve activation respectively. Subsequently, the LF/HF power ratio has been proposed as a compact indicator of sympato-vagal balance, however, it has been noted that the correspondence of sympathetic and vagal activation to the LF and HF band powers might not be so straightforward, but a more complex interaction, and that the proportionalisation of the two band-powers further contracts the sympathetic, parasympathetic and possibly other underlying independent factors into a single measure [Billman et al. 2013].

As many other electrophysiological signals (local field potentials, electrocorticograms, electroencephalograms, magnetoencephalograms), the RR-interval time series also contain an aperiodic component besides oscillations. The approach of separating the oscillatory and aperiodic components of the spectrum has gained popularity in the field of electroencephalography. [Donoghue et al. 2020ab, Wen et al. 2015] as changes in the aperiodic component carry meaningful information on brain states [Colombo et al. 2019, Schneider et al. 2022, Maschke et al. 2023]. Thus, untangling the oscillatory and aperiodic contributions could be important in the case of HRV signals as well, as they might reflect different underlying heartbeat regulation mechanisms, which traditional band-based approaches (like calculating the total, LF or HF powers) might confuse.

It has been revealed that heart rate fluctuations do not only conform to a power-law PSD characterized by physiologically and cardiologically relevant parameters (Kobayashi and Musha, 1982; Bigger et al., 1996), but also uncover a fractal temporal structure of the underlying tachogram [Yamamoto et al. 1994]. Moreover, the known multifractal nature of RR-interval time series appears to be an essential property of healthy heart functioning [Peng et al., 1995, Ivanov et al. 1999], and possibly a consequence of modulatory mechanisms spanning several temporal scales [Monfredi & Lakatta, 2019], which further indicates that a multi-exponent description of the RR-interval spectra is necessary.

Our aim is to uncover the fundamental aspects of the sleep-related multi-exponent human RR-interval spectra. Our study presents effects of aging and sex on heart rate variability during sleep in a healthy population, using a novel, parametric description of RR-interval power spectra, through the adoption of a broken power-law model characterizing the fractal component with two spectral slopes and intercepts separated by a custom breaking-point, in addition to the parametrization of the LF and HF band peaks. Furthermore, we investigate the relationship between fractal and oscillatory spectral parameters to classical HRV measures and sleep macrostructure indicators. Based on the evidence supporting the multifractality of the time series representing healthy human RR sequences [Peng et al., 1995, Ivanov et al., 1999], as well as the relationship between the fractal scalings in the time and the frequency domain, we hypothesize that heartbeat dynamics during sleep follow a broken power law-type spectra, with two exponents and a breaking point. Moreover, we hypothesize that oscillations corresponding to Mayer waves (∼0.1 Hz) and to respiratory sinus arrhythmia (∼0.2 Hz) are forming non-fractal, periodic activities, which lead to spectral peaks at respective frequencies. Furthermore, we hypothesize that known age- and sex differences in heart rate dynamics are reflected in the parameters describing multifractal and oscillatory activities of the human RR-spectra during sleep. Last, but not least, we hypothesize that objective, polysomnography-based differences in sleep quality are reflected in fractal and oscillatory measures of RR time series spectra.

## Methods

Full-night ECG recordings were extracted from an already existing polysomnography database of 251 healthy humans from a wide age range (4–69 years), 122 females (Budapest-Munich database). Subjects had given their written consent for inclusion in the database and the multilaboratory data collection procedures had been approved by corresponding local ethical committees, all respecting the Declaration of Helsinki (Bódizs et al., 2017). The duration of records varied between 212 and 729 minutes (mean: 513.44 minutes, SD:63.06 minutes). All records contained one bipolar ECG derivation according to the polysomnography standards. R-peaks were automatically detected, then the evenly resampled (4 Hz sampling rate) RR-interval time series constructed using the Kubios HRV software [Travainen et al. 2014]. Missing segments due to artefact removal in the series were replaced with values linearly interpolated between the last and first values before and after the gaps. Recordings that contained more than 10% missing values were excluded from the analysis, leaving 215 high-quality RR-interval series (103 females, age range: 4–69, mean: 26.44 years). The fractal and oscillatory PSD of the whole-night RR-interval time series were calculated using the Irregular-Resampling Auto-Spectral Analysis (IRASA) method [Wen et al. 2015], in the frequency range of 0.001–0.5 Hz, using a moving window of 4096 seconds.

### Parametrization of the fractal component

Transforming both the power and frequency axes of the fractal component of the PSD into logarithmic space, it can be noted that the relation between power and frequency is described by a broken power-law [see Fig. 1. left]. After uniform resampling of the PSD in the logarithmic frequency domain, piecewise linear regression [Jekel et al. 2019] was applied to the fractal component, which automatically locates a breaking point and determines slope and intercept values of two linear functions for the lower and higher frequency domains. To confirm that the broken power-law model was more appropriate than the single slope description, the Akaike information criterion was calculated for simple linear and two-slope piecewise regressions in the log-log domain. Lower Akaike information criterion values indicate better model fits. As the metric is derived from the residual sum of squares but includes a penalty term for the number of model parameters, thus overfitting and unnecessary model complexity can be avoided, in contrast to simple goodness-of-fit metrics like the R^2^ coefficient of determination. [Gkioulekas, I., & Papageorgiou, L. G. 2018]

**Figure 1.**
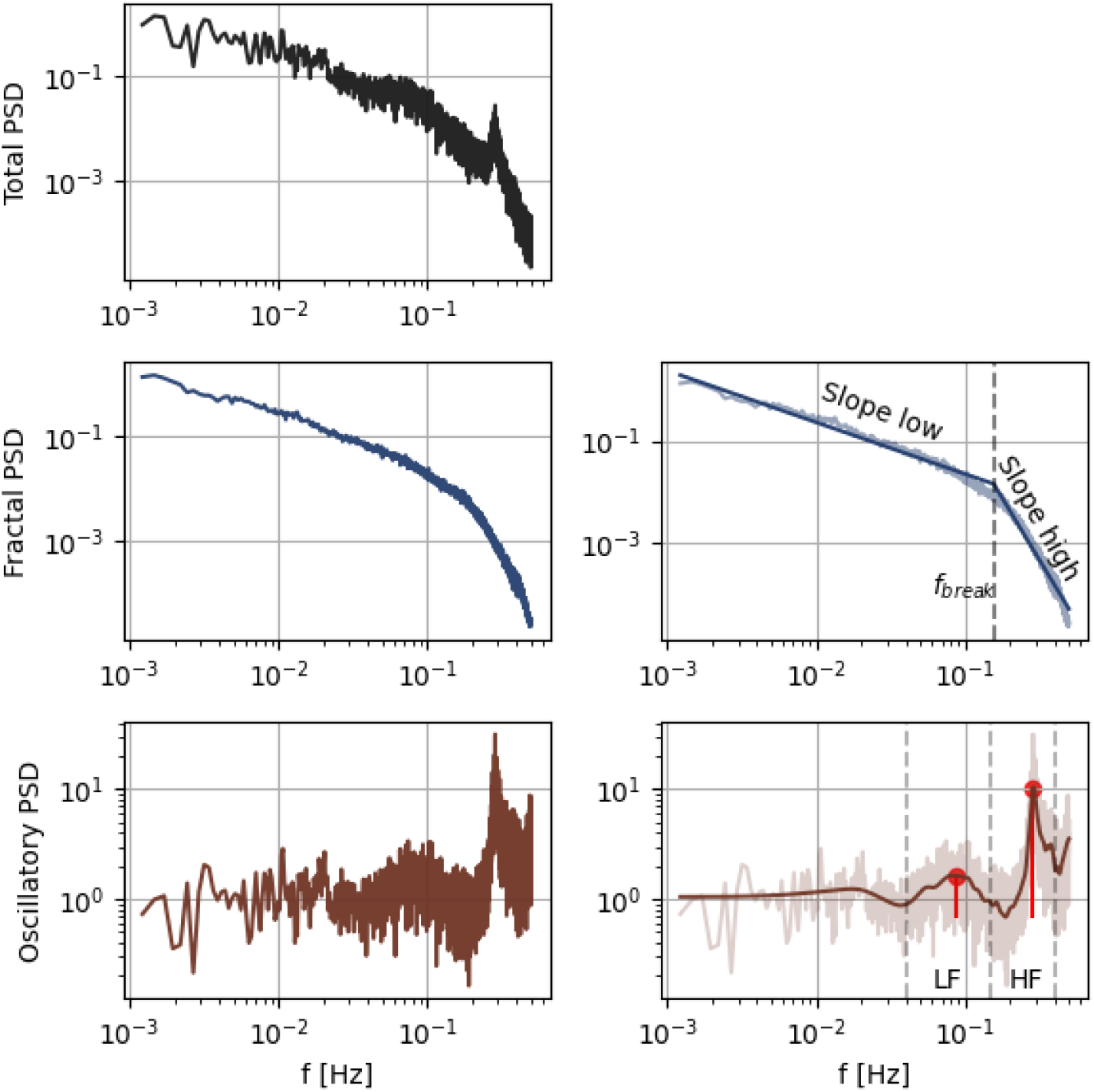
Total RR-interval power-density spectra (top left), fractal component separated by IRASA (middle left) and the fitted piecewise linear function (middle right). The oscillatory component (bottom left) and peaks detected in the LF and HF bands after gaussian smoothing (bottom right).

**Figure 2.**
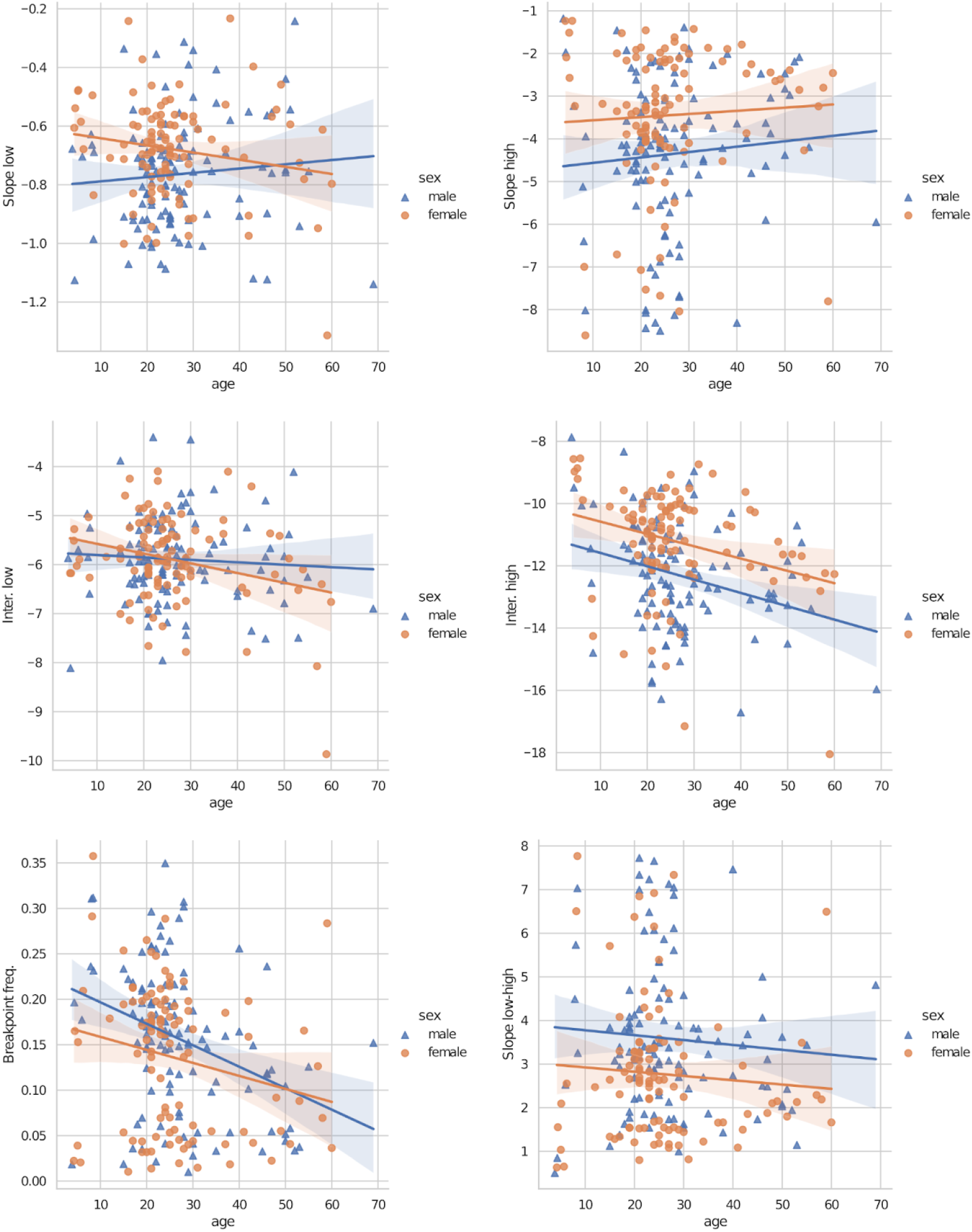
ANCOVA plots of the fractal parameters, significant effects of age were found in the high domain intercept (F(1,212)=19.107, p<0.001, η^2^_p_=0.083, middle right) and breaking point frequency (F(1,206)=18.08, p<0.001, η^2^_p_=0.022, bottom left), furthermore effects of sex in both low domain slope (F(1, 212)=11.64, p<0.001, η^2^_p_=0.052, top left) and high domain slope (F(1, 212)=15.51, p<0.001, η^2^_p_=0.068, top right), high domain intercept (F(1, 212)=22.988, p<0.001, η^2^_p_=0.098, middle right) and slope difference (F(1,212)=13.899, p<0.001, η^2^_p_=0.062, bottom right). (N=215 subjects, 122 females)

**Figure 3.**
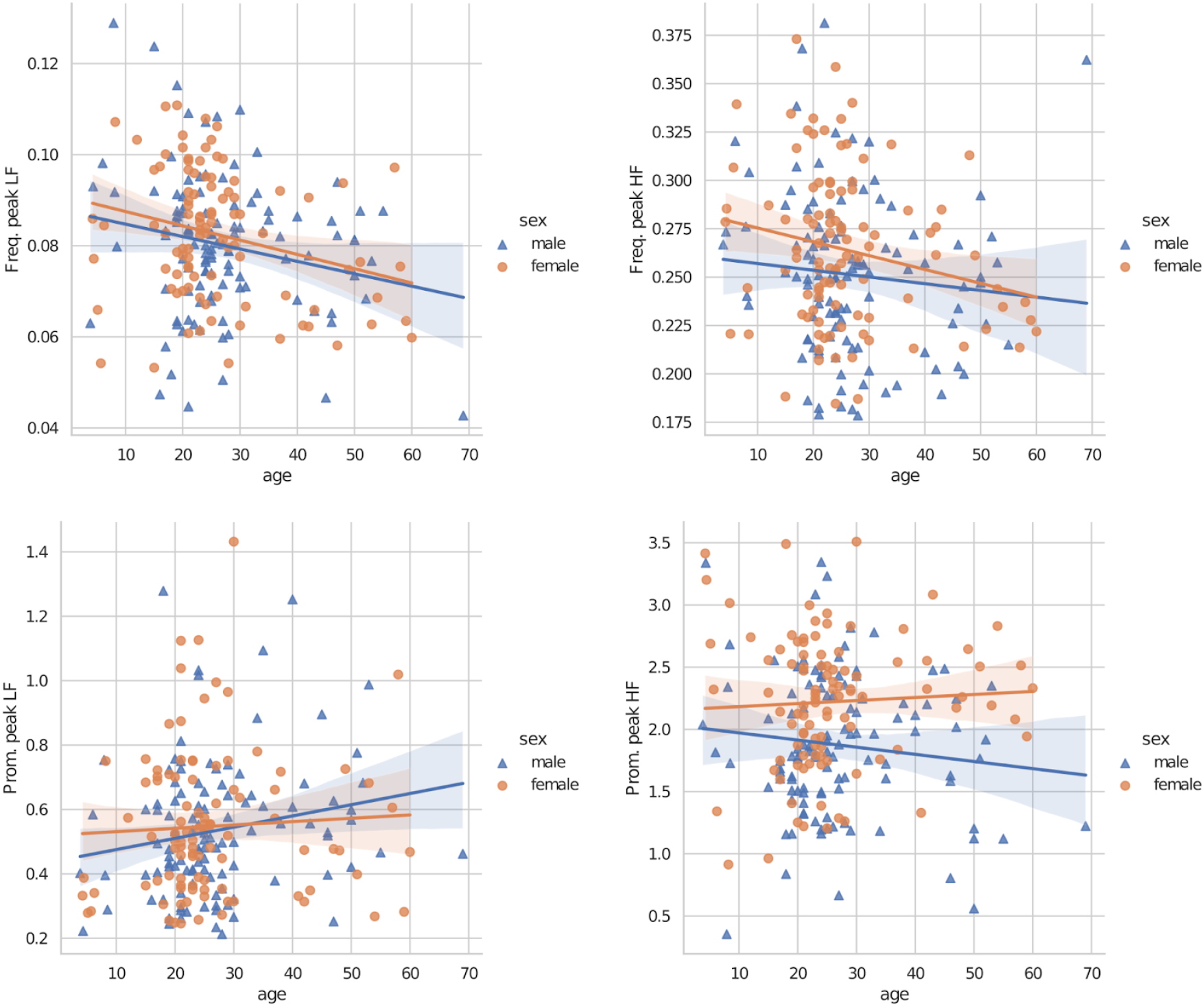
ANCOVA plot of oscillatory parameters, significant effects of age in the LF peak frequency (F(1,206)=11.502, p<0.001, η^2^_p_ =0.053, top left) and sex in the HF peak prominence (F(1, 212)=22.988, p<0.001, η^2^_p_=0.098, bottom right). (N=215 subjects, 122 females)

### Parametrization of the oscillatory component

The oscillatory PSD was established by subtracting the fractal component from the total PSD in the log-log space, then smoothed with a gaussian filter (the standard deviation of the gauss kernel was 20 samples). The frequencies and prominences of the dominant peaks were extracted via a peak detection algorithm [Virtanen et al. 2020] in the low frequency (LF) 0.04–0.15 Hz and high frequency (HF) 0.15–0.4 Hz bands commonly used in HRV analyses.

After extracting the spectral parameters, ANCOVA analysis was carried out for each parameter as the dependent variable with sex as a fixed factor and age as a covariate.

Using manually scored and validated hypnograms from the polysomnography database, macrostructure indicators were calculated. The indicators included total sleep time (TST, duration of all sleep stages in the recording), sleep efficiency (SE, ratio of TST compared to the duration of the recording), wake after sleep onset (WASO, total time in wake state after the first N2 and last sleep stage) and the percentage of each sleep stage (compared to the time between the first N2 sleep and last sleep stage), then were Pearson correlated with the overnight spectral parameters of heart rate, correlations were considered significant after Hom-Bonferroni correction (p_k_<0.005/(m-k+1); k=1,…,m; m=63;).

Classical HRV indicators were also calculated, the HRV index (total spectral power), then LF and HF power, finally the LF/HF ratio. Multiple linear regression was carried out for each classical HRV marker as the dependent variable and groups of spectral parameters as predictors.

## Results

### Multifractality of human heart rate dynamics during sleep

We revealed a lower Akaike Information Criterion value for the two-slope models in all 215 cases, suggesting a better model fit with two spectral slopes as compared to the classical one exponent solutions reported in the literature on the power law scaling of the Fourier transform-derived periodogram of human RR series.

The overall descriptives of the spectral parameters show that the spectral slope was generally flatter in the low frequency domain (M=-0.723) than in the high domain (M=-3.916), the difference of the two slopes being positive in all cases, besides that the breaking point frequency varies in a wide range (range=0.010–0.358 Hz). The coefficients of determination (r-squared) for the piecewise regression indicated adequate fits for the fractal PSD component (mean=0.983, range=0.944–0.997), for more descriptives see Table 1.

**Table 1.** General descriptives of the fractal spectral parameters.

| Descriptive Statistics |  |  |  |  |  |  |  |
| --- | --- | --- | --- | --- | --- | --- | --- |
|  | slope1 | slope2 | slope_diff | inter1 | inter2 | fbreak | rsqr |
| Valid | 215 | 215 | 215 | 215 | 215 | 215 | 215 |
| Missing | 0 | 0 | 0 | 0 | 0 | 0 | 0 |
| Mean | -0.723 | -3.916 | 3.193 | -5.889 | -11.783 | 0.147 | 0.983 |
| Std. Deviation | 0.182 | 1.748 | 1.673 | 0.862 | 1.769 | 0.080 | 0.008 |
| Minimum | -1.314 | -8.605 | 0.501 | -9.869 | -18.051 | 0.010 | 0.944 |
| Maximum | -0.233 | -1.179 | 7.769 | -3.402 | -7.870 | 0.358 | 0.997 |

Peaks were identified in almost all oscillatory spectra in the LF and HF domain (with 6 and 2 exceptions respectively), HF peaks being more prominent as expected, see Table 2.

**Table 2.** Descriptive statistics of the fitted oscillatory parameters.

| Descriptive Statistics |  |  |  |  |
| --- | --- | --- | --- | --- |
|  | fpeak_lf | fpeak_hf | prom_lf | prom_hf |
| Valid | 209 | 213 | 209 | 213 |
| Missing | 6 | 2 | 6 | 2 |
| Mean | 0.081 | 0.257 | 0.541 | 2.039 |
| Std. Deviation | 0.015 | 0.041 | 0.218 | 0.582 |
| Minimum | 0.043 | 0.178 | 0.213 | 0.354 |
| Maximum | 0.129 | 0.381 | 1.432 | 3.509 |

### Age effects

A high frequency domain intercept decrease was observed with aging [F(1,212)=19.107, p<0.001, η^2^_p_ =0.083], while the breaking point frequency also showed a decreasing tendency [F(1,206)=18.08, p<0.001, η^2^_p_ =0.022]. Oscillatory peak frequency exhibited slowing as age advances in the LF domain [F(1,206)=11.502, p<0.001, η^2^_p_ =0.053].

### Sex effects

Significant effects of sex were found both in the low domain slope [M_female_=-0.680, SD_female_=0.165, M_male_=-0.764, SD_male_=0.189, F(1, 212)=11.64, p<0.001, η^2^_p_=0.052] and high domain slope [M_female_=-3.448, SD_female_=1.161, M_male_=-4.347, SD_male_=1.760, F(1, 212)=15.51, p<0.001, η^2^_p_=0.068] with increased values indicating flatter spectra in females. The high domain intercept showed a consistent increase in female subjects [F(1, 212)=22.988, p<0.001, η^2^_p_=0.098] as well. Furthermore, the slope difference between the low and high domain was significantly higher in males [F(1,212)=13.899, p<0.001, η^2^_p_=0.062]. Considering the oscillatory component, the HF peak was more prominent in females [F(1,210)=20.451, p<0.001, η^2^_p_=0.089].

### Relationship to sleep macrostructure indicators

In the following, the effect of sleep structure on the spectral parameters was examined, WASO correlated negatively with the spectral intercept in both low [r=-0.290] and high [r=-0.270] domains, furthermore N1 percentage was also anticorrelated with the high domain intercept [r=-0.298], while R percentage showed positive association with the low domain intercept [r=0.303], see Figure 4.

**Figure 4.**
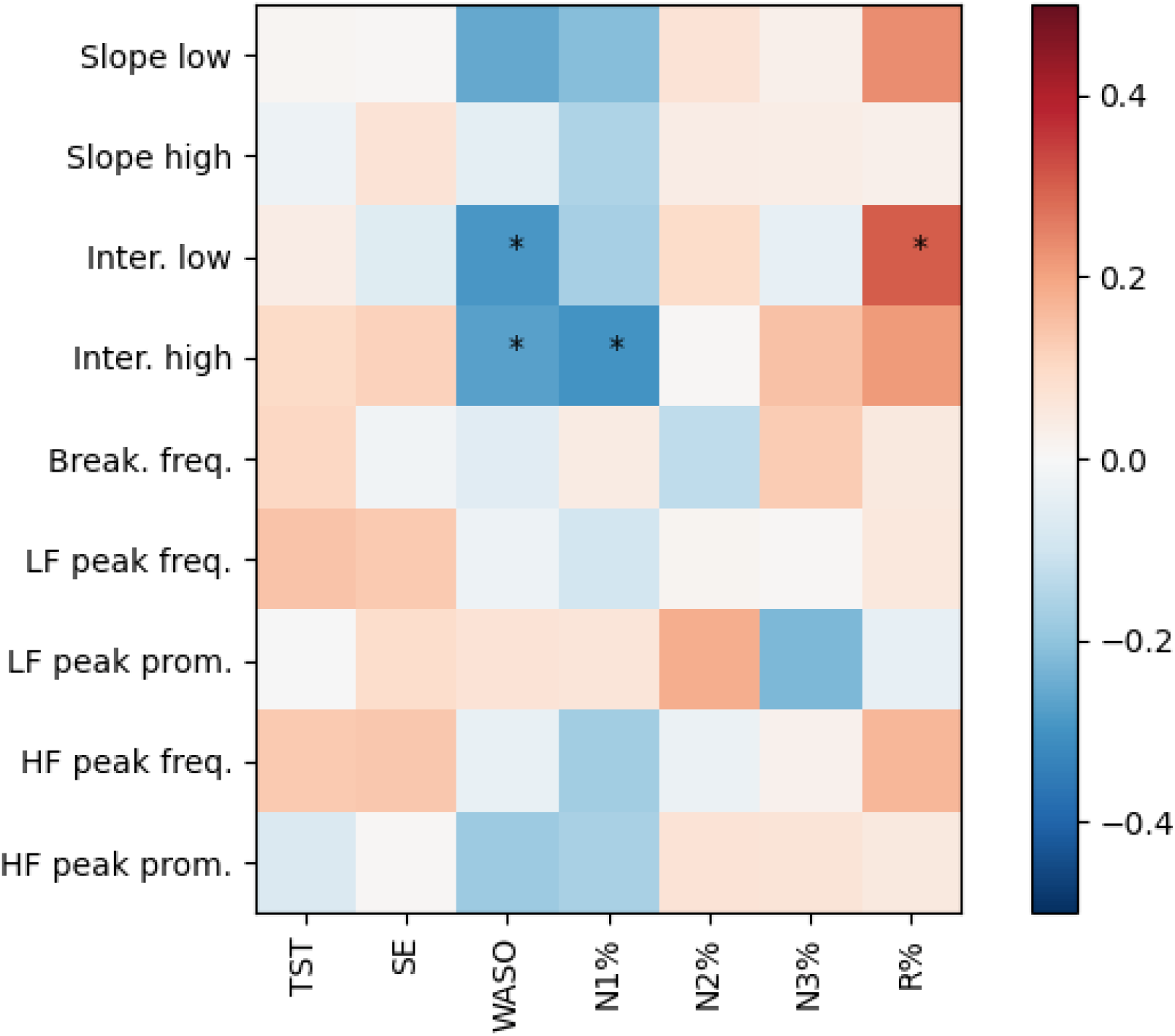
Pearson correlation matrix of the spectral parameters and the sleep macrostructure indicators, significant relationships marked with “^*^“, Bonferroni-Holm corrected (ɑ=0.005). (TST - total sleep time, SE - sleep efficiency, WASO - wake after sleep onset, and sleep stage duration percentages).

### Relationship to classical HRV markers

In order to assess which of our model parameters account for the individual differences in classical HRV measures, a multiple linear regression was implemented by considering each classical HRV marker as a dependent variable, and the fractal, oscillatory, then all spectral parameters as predictors. In all cases a substantial amount of the variance in the traditional, band-limited spectral variables were explained by changes in the fractal parameters, see Table 3.

**Table 3.**
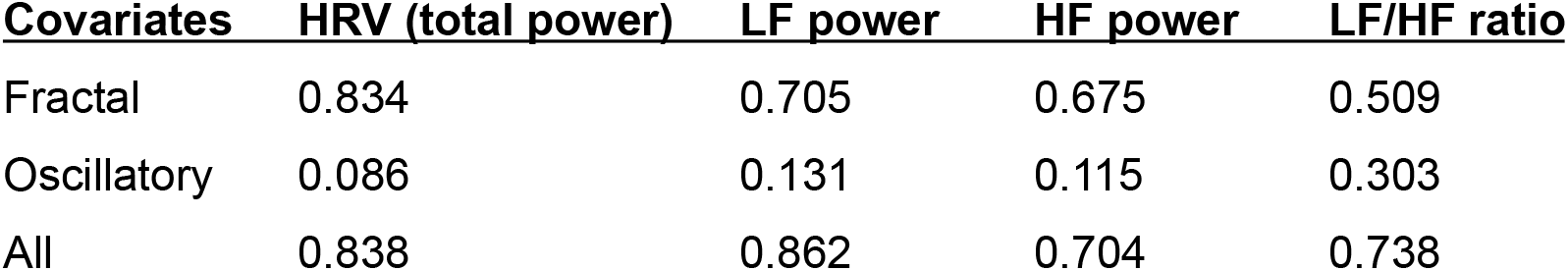
Variance of classical HRV indicators (columns) explained by groups of spectral parameters (rows), R^2^ values of multiple linear regression tests. The spectral parameters were grouped as Fractal: low and high domain slope, low domain intercept, breaking point frequency; Oscillatory: peak frequency and prominence in the LF and HF bands; All: both fractal and oscillatory parameters.

## Discussion

Fractal spectra are ubiquitous in natural time series, including physiological rhythms and fluctuations. Former studies described the power law scaling of heart rate fluctuation spectra (Kobayashi and Musha, 1982), revealing the physiological and cardiological relevance of the spectral parameters (Biggers et al., 1996). Moreover, sleep-associated heart rate was shown to characterize sleep quality and health (Stein and Pu, 2012). Last, but not least, indirect evidence suggests the bi-fractal nature of human heart rate, characterized by different parameters in different frequency ranges (Peng et al., 1995, Ivanov et al., 1999). Here we implemented a novel approach in determining sleep-related heart rate variation spectra with the aim of explicitly targeting a broken power law model-type description of the periodograms reflecting cardiac autonomic modulation in humans. As the two-slope model provided a more accurate description of the fractal spectra (also reflected by reduced Akaike information criteria), it evidenced of the presence of a significant breaking point in the power-spectrum, which also carried physiological significance. Findings replicate and specify some of the assertions on the relatedness of sleep quality and the autonomic modulation of heart rate.

The piecewise linear regression approach applied to the fractal spectra of equidistantly resampled RR-series revealed novel spectral parameters of heart rate variability, which are related to, but not equivalent with the well-known frequency domain HRV measures of the band-limited type (eg. total, LF, HF, power or LF/HF ratio). Consequently, some general comparisons can be made with the literature to interpret and confirm our results based on the aforementioned novel parameters.

Our finding of a significant intercept reduction in the high frequency domain with aging coheres with the age-related nocturnal HRV attenuation indicating decreased adaptability [Reardon et al. 1996, Kentta et al., 2020]. Furthermore, published results indicate that age primarily affects the higher end of the spectrum [Olivieri et al. 2024], a finding that parallels the age-relatedness of the respective fractal parameters of our current study. Taken together with the new observation that the breaking point frequency also showed age-related reduction, and that there were no significant changes in the spectral slopes or peak prominences with age, we propose that the reduction of the total HRV power with aging can be explained by a shift in the broken power-law towards lower frequencies.

Looking at known sex differences in HRV [Koenig et al. 2016], the attenuation of total HRV power in females compared to males is not obviously reflected in our results, however greater HF power supports our findings of increased high domain intercepts and HF peak prominences in females, furthermore increased LF/HF ratios are aligned with the flatter spectral slopes in both domains in our model.

As the dynamics of sleep and cardiac regulation are tied together by modulations in the ANS, HRV is also an indicator of sleep quality, being reduced in improper sleep [Sajjadieh et al. 2020], [deEstrela et al. 2020][Schlagintweit et al., 2023], which confirms our results that more fragmented sleep is associated with an overall decrease in HRV power, as both the low and high domain intercepts showed significant anticorrelation with WASO duration. Furthermore, REM sleep is known to increase sympathetic tone, which is considered to be reflected by LF power increase [Elsenbruch et al. 1999], overlapping with the positive correlation between our low domain intercept and REM sleep percentage, note that longer episodes of REM sleep can also be associated with quality sleep, as REM becomes more prominent with the dissipation of homeostatic sleep pressure. Consequently, the link between sleep and cardiac regulation could be utilized. On the one hand, while within-stage HRV measurements are known to be stable indicators [Kerkering et al., 2022], overnight variability measures are correlated with aspects of sleep structure, allowing for the assessment of sleep quality from heart rate data. On the other hand, in the future polysomnographic recordings might also provide data for automated cardiovascular health screenings.

Distinct oscillations were found in almost all cases in the LF and HF bands in the whitened oscillatory spectra, in addition the mean breaking point frequency in the fractal component was close to the 0.15 Hz boundary separating the two bands, which indicates the validity of splitting the two time scales, however the variance and the discovered age effect on the breaking point frequency advises against setting a fixed boundary for the low and high frequency bands. The proposed method is able to describe the complete frequency-domain landscape of HRV in a single model, in contrast to traditional metrics that were based on fixed frequency bands, furthermore can differentiate between oscillatory and fractal changes. In the present study only the LF and HF bands were covered due to the limited duration of the ECG recordings, however the model is easily extendable to the VLF or ULF domains, given proper data. Most of the power variance was explained by aperiodic spectral parameters, while age and sex effects were attributed also to fractal parameters in greater numbers, so we propose that the changes occurring in the fractal component of the HRV spectra dominate these effects, in contrast to oscillations, and have to be considered in further studies.

## References

Alrosan, A. Z., Heilat, G. B., Alrosan, K., Aleikish, A. A., Rabbaa, A. N., Shakhatreh, A. M., Alshalout, E. M., & Al Momany, E. M. A. (2024). Autonomic brain functioning and age-related health concerns. Current Research in Physiology, 7, 100123. 10.1016/j.crphys.2024.100123

Bigger, J. T., Jr, Steinman, R.C., Rolnitzky, L. M., Fleiss, J. L., Albrecht, P., & Cohen, R. J. (1996). Power Law Behavior of RR-Interval Variability in Healthy Middle-Aged Persons, Patients With Recent Acute Myocardial Infarction, and Patients With Heart Transplants. Circulation, 93(12), 2142–2151. 10.1161/01.cir.93.12.2142

Billman, G. E. (2013). The LF/HF ratio does not accurately measure cardiac sympatho-vagal balance. Frontiers in Physiology, 4. 10.3389/fphys.2013.00026

Bódizs, R., Gombos, F., Ujma, P. P., Szakadát, S., Sándor, P., Simor, P., Pótári, A., Konrad, B. N., Genzel, L., Steiger, A., Dresler, M., & Kovács, I. (2017). The hemispheric lateralization of sleep spindles in humans. Sleep Spindles & Cortical Up States, 1(1), 42–54. 10.1556/2053.01.2017.002

ChuDuc, H., NguyenPhan, K., & NguyenViet, D. (2013). A Review of Heart Rate Variability and its Applications. APCBEE Procedia, 7, 80–85. 10.1016/j.apcbee.2013.08.016

Colombo, M. A., Napolitani, M., Boly, M., Gosseries, O., Casarotto, S., Rosanova, M., Brichant, J.-F., Boveroux, P., Rex, S., Laureys, S., Massimini, M., Chieregato, A., & Sarasso, S. (2019). The spectral exponent of the resting EEG indexes the presence of consciousness during unresponsiveness induced by propofol, xenon, and ketamine. NeuroImage, 189, 631–644. 10.1016/j.neuroimage.2019.01.024

Donoghue, T., Haller, M., Peterson, E. J., Varma, P., Sebastian, P., Gao, R., Noto, T., Lara, A. H., Wallis, J. D., Knight, R. T., Shestyuk, A., & Voytek, B. (2020a). Parameterizing neural power spectra into periodic and aperiodic components. Nature Neuroscience, 23(12), 1655–1665. 10.1038/s41593-020-00744-x

Donoghue, T., Dominguez, J., & Voytek, B. (2020b). Electrophysiological Frequency Band Ratio Measures Conflate Periodic and Aperiodic Neural Activity. Eneuro, 7(6), ENEURO.0192-20.2020. 10.1523/eneuro.0192-20.2020

da Estrela, C., McGrath, J., Booij, L., & Gouin, J.-P. (2020). Heart Rate Variability, Sleep Quality, and Depression in the Context of Chronic Stress. Annals of Behavioral Medicine, 55(2), 155–164. 10.1093/abm/kaaa039

Elsenbruch, S., Harnish, M. J., & Orr, W. C. (1999). Heart Rate Variability During Waking and Sleep in Healthy Males and Females. Sleep, 22(8), 1067–1071. 10.1093/sleep/22.8.1067

Gkioulekas, I., & Papageorgiou, L. G. (2018). Piecewise Regression through the Akaike Information Criterion using Mathematical Programming. IFAC-PapersOnLine, 51(15), 730–735. 10.1016/j.ifacol.2018.09.168

Ivanov, P. Ch., Amaral, L. A. N., Goldberger, A. L., Havlin, S., Rosenblum, M. G., Struzik, Z. R., & Stanley, H. E. (1999). Multifractality in human heartbeat dynamics. Nature, 399(6735), 461–465. 10.1038/20924

Jekel, Charles F. and Venter, Gerhard (2019) PWLF: A Python Library for Fitting 1D Continuous Piecewise Linear Functions https://github.com/cjekel/piecewise_linear_fit_py

Kentta, T. V., Karsikas, M., Rantanen, A., Kinnunen, H., & Koskimaki, H. (2020). Age, sex and underlying heart rate as determinants of nocturnal heart rate variability among wearable smart ring users. European Heart Journal, 41(Supplement_2). 10.1093/ehjci/ehaa946.3463

Kerkering, E. M., Greenlund, I. M., Bigalke, J. A., Migliaccio, G. C. L., Smoot, C. A., & Carter, J. R. (2022). Reliability of heart rate variability during stable and disrupted polysomnographic sleep. American Journal of Physiology-Heart and Circulatory Physiology, 323(1), H16–H23. 10.1152/ajpheart.00143.2022

Kobayashi, M., & Musha, T. (1982). 1/f Fluctuation of Heartbeat Period. IEEE Transactions on Biomedical Engineering, BME-29(6), 456–457. 10.1109/tbme.1982.324972

Koenig, J., & Thayer, J. F. (2016). Sex differences in healthy human heart rate variability: A meta-analysis. Neuroscience & Biobehavioral Reviews, 64, 288–310. 10.1016/j.neubiorev.2016.03.007

Maschke, C., Duclos, C., Owen, A. M., Jerbi, K., & Blain-Moraes, S. (2023). Aperiodic brain activity and response to anesthesia vary in disorders of consciousness. NeuroImage, 275, 120154. 10.1016/j.neuroimage.2023.120154

Monfredi, O., & Lakatta, E. G. (2019). Complexities in cardiovascular rhythmicity: perspectives on circadian normality, ageing and disease. Cardiovascular Research, 115(11), 1576–1595. 10.1093/cvr/cvz112

Nagai, M., Hoshide, S., & Kario, K. (2010). Sleep Duration as a Risk Factor for Cardiovascular Disease-a Review of the Recent Literature. Current Cardiology Reviews, 6(1), 54–61. 10.2174/157340310790231635

Olivieri, F., Biscetti, L., Pimpini, L., Pelliccioni, G., Sabbatinelli, J., & Giunta, S. (2024). Heart rate variability and autonomic nervous system imbalance: Potential biomarkers and detectable hallmarks of aging and inflammaging. Ageing Research Reviews, 101, 102521. 10.1016/j.arr.2024.102521

Peng, C.-K., Havlin, S., Stanley, H. E., & Goldberger, A. L. (1995). Quantification of scaling exponents and crossover phenomena in nonstationary heartbeat time series. Chaos: An Interdisciplinary Journal of Nonlinear Science, 5(1), 82–87. 10.1063/1.166141

Reardon, M., & Malik, M. (1996). Changes in Heart Rate Variability with Age. Pacing and Clinical Electrophysiology, 19(11), 1863–1866. 10.1111/j.1540-8159.1996.tb03241.x

Sajjadieh A, Shahsavari A, Safaei A, Penzel T, Schoebel C, Fietze I, Mozafarian N, Amra B, Kelishadi R. The Association of Sleep Duration and Quality with Heart Rate Variability and Blood Pressure. Tanaffos. 2020 Nov;19(2):135–143. PMID: 33262801; PMCID: PMC7680518.

Schlagintweit, J., Laharnar, N., Glos, M., Zemann, M., Demin, A. V., Lederer, K., Penzel, T., & Fietze, I. (2023). Effects of sleep fragmentation and partial sleep restriction on heart rate variability during night. Scientific Reports, 13(1). 10.1038/s41598-023-33013-5

Schneider, B., Szalárdy, O., Ujma, P. P., Simor, P., Gombos, F., Kovács, I., Dresler, M., & Bódizs, R. (2022). Scale-free and oscillatory spectral measures of sleep stages in humans. Frontiers in Neuroinformatics, 16. 10.3389/fninf.2022.989262

Stein, P. K., & Pu, Y. (2012). Heart rate variability, sleep and sleep disorders. Sleep Medicine Reviews, 16(1), 47–66. 10.1016/j.smrv.2011.02.005

Tarvainen, M. P., Niskanen, J.-P., Lipponen, J. A., Ranta-aho, P. O., & Karjalainen, P. A. (2014). Kubios HRV – Heart rate variability analysis software. Computer Methods and Programs in Biomedicine, 113(1), 210–220. 10.1016/j.cmpb.2013.07.024

Tobaldini, E., Nobili, L., Strada, S., Casali, K. R., Braghiroli, A., & Montano, N. (2013). Heart rate variability in normal and pathological sleep. Frontiers in Physiology, 4. 10.3389/fphys.2013.00294

Virtanen, P., Gommers, R., Oliphant, T. E., Haberland, M., Reddy, T., Cournapeau, D., Burovski, E., Peterson, P., Weckesser, W., Bright, J., van der Walt, S. J., Brett, M., Wilson, J., Millman, K. J., Mayorov, N., Nelson, A. R. J., Jones, E., Kern, R., Larson, E., … Vázquez-Baeza, Y. (2020). SciPy 1.0: fundamental algorithms for scientific computing in Python. Nature Methods, 17(3), 261–272. 10.1038/s41592-019-0686-2

Wen, H., & Liu, Z. (2015). Separating Fractal and Oscillatory Components in the Power Spectrum of Neurophysiological Signal. Brain Topography, 29(1), 13–26. 10.1007/s10548-015-0448-0

Yamamoto, Y., & Hughson, R. L. (1994). On the fractal nature of heart rate variability in humans: effects of data length and beta-adrenergic blockade. American Journal of Physiology-Regulatory, Integrative and Comparative Physiology, 266(1), R40–R49. 10.1152/ajpregu.1994.266.1.r40

